# Persistent repetition of brain networks in *robust Lesion Network Mapping:* A reply to *Cui et al*

**DOI:** 10.64898/2026.11.17.767082

**Authors:** Ilan Libedinsky, Sebastian Quiroz Monnens, Marius Gruber, Jonathan Repple, Luca Cocchi, Martijn P. van den Heuvel

## Abstract

Lesion Network Mapping (LNM) include a series of methods used for identifying disease brain circuits. An investigation into the method’s methodological roots showed persistent repetition among derived networks. Recently, “robust” LNM (rLNM) is proposed as an update to the analysis procedures. By adding new statistical steps and layers, rLNM regresses out a common cross-disorder connectivity factor with the claim to reduce repetition and improve network specificity. However, rLNM remains to rely on projecting coordinates and lesion locations onto a single normative functional connectome *C*. Statistical procedures such as specificity testing or rLNM cross-symptom regression may alter the resulting maps, but they do not change the fundamental dependence on *C*. Here, we discuss the rLNM variant in the context of the lesion set *M* and the connectivity matrix *C*. We show that this new variant effectively only adjusts the lesion matrix of group A relative to groups B and C, replacing *M*_*A*_ to *M′*_*A*_ *= (2M*_*A*_ − *M*_*B*_ − *M*_*C*_*)/3*. This leaves the core procedure to *LNM ∼ M′*_*A*_*C* and its outcomes subject to the same connectome-driven repetition when tested across multiple conditions. We empirically evaluate the rLNM approach across 67 disorders and their 2-group and 3-group combinations. We show that the new procedure unfortunately leaves the characteristic repetition among networks, producing overlapping circuits for unrelated clinical conditions.

## Main text

Lesion Network Mapping (LNM)^1-4^ is a neuroimaging method that projects coordinates and lesion locations onto a normative functional connectome to identify brain circuits associated with clinical symptoms^5^. The method is also known under various other formulations such as coordinate network mapping^5^, atrophy network mapping^6^, activation network mapping^7^, convergent connectivity mapping^8, 9^, therapeutic network mapping^10^, glioma network mapping^11^, and others^7, 12, 13^.

An investigative study^14^ into the methodological steps of LNM raised concerns about high repetition and low anatomical specificity among reported LNM networks, with circuits proposed for distinct psychiatric and neurological conditions overlapping at |r| > 0.6-0.8^15-18^. This overlap is much higher than typically seen in neuroimaging studies, approaching an empirical reliability ceiling for LNM and functional connectivity estimates^19, 20^.

This repetition is not biological, but methodological in origin^14^: Beneath the framework’s complexity, LNM involves a linear matrix operation, *MC*, with lesions *M* projected onto a single normative healthy brain functional connectome matrix *C*^*14*^. Subsequent discussion^19, 21^ clarified that repeatedly sampling the same low-dimensional matrix by *MC* can readily present standard brain connectivity patterns as disease circuits when tested within and across conditions and contrasts. This repetition remains when statistical steps such as specificity testing and null-models are applied, as they are mathematically only updates of the lesion matrix *M*—e.g. *M = (M*_*A*_ *− M*_*B*_) in specificity testing^22^ or *M = (M − 1/R)* in random-lesion null models^23^—leaving the core procedure unchanged to a *MC* operation.

Cui and colleagues^24^ propose a new null-model-based extension to the LNM framework, building on null-models for LNM. Partly acknowledging the imprint of a repetitive core pattern (“degree”), the authors propose a series of steps collectively termed *robust LNM* (*rLNM*), with the goal of regressing out the common factor to derive symptom-specific networks^24^.

The presented rationale for this approach is that the reported overlap among LNM networks is perceived as a consequence of the “degree” or “hub architecture”^21, 25, 26^ of the normative connectome, arguing that statistical correction of a labelled “connectome-driven bias”^*24*^ will lead to brain circuits with anatomical specificity across conditions. The original investigative study already discussed this narrow interpretation of convergence on brain hubs^14^: The broader concern is that LNM operations inherently project lesions onto a single low-rank normative matrix, leaving little room for varying outcomes. LNM and its variants, such as coordinate network mapping, remain to project a lesion matrix *M*—or updated versions *M* such as *(M − 1/R)*, or as we will derive below in *rLNM ∼ (2M*_*A*_ *− M*_*B*_ *− M*_*C*_) when testing three groups—to the same normative connectivity matrix *C*. If the procedure remains an operation of *MC*, the possible space of outputs necessarily remains constrained to the patterns embedded within *C*^19, 27, 28^.

Null models are valuable tools in network neuroscience^29^, and the proposed rLNM method combines several useful concepts and statistical steps, warranting examination and discussion. This is particularly relevant because researchers are rapidly adopting such null models^30-32^ as possible solutions to the limitation of LNM. Below, we discuss the steps of the new rLNM variant in the context of *M* and *C*: We revisit the *“null-based sensitivity step”* (proposed to replace LNM sensitivity testing) and show that, when expressed in its mathematically equivalent form, it only reformulates standard specificity testing and earlier proposed random-coordinate null-models^1, 9^. We next assess the second multi-step procedure *“specificity testing with cross-map regression”* of rLNM and derive that, when studied in context of *M* and *C*, it effectively also only updates matrix *M* for group A by subtracting those of group B and group C, leaving the series of steps to a (*M*_*A*_ *− M*_*BC*_*)C* operation, and with that retaining the linear dependence on matrix *C*. We empirically evaluate the rLNM approach across 67 coordinate and lesion sets and their 2 and 3-group combinations and show that rLNM maps continue to show the characteristic repetition of networks seen in LNM.

## Results

### Robust LNM: Null-model controlled sensitivity

The standard LNM procedure, can be written as *LNM = ∑(MC)*^14^, or *MC* for short, with *M* a lesion-size and set-size normalised matrix containing the lesions or coordinates of the symptom of interest and *C* the standard normative connectivity matrix^14^. In rLNM, Cui et al. propose modifying the standard LNM sensitivity step^24^ by contrasting the fc maps of the lesions of interest with those derived from random lesions/coordinates^24^. The method compares the standard sensitivity LNM map against a null distribution of LNM maps obtained from e.g. 5,000 random lesion sets, with the output map labelled as the null-controlled sensitivity map^24^. This approach is comparable to proposed null models implementing spin-randomised lesions^14^ or random coordinates^1, 9^.

In more detail, the random-lesion null-model procedure consists of (1) generating the observed LNM map, *LNMobs*; (2) repeatedly sampling an equal number of random lesions or coordinates from the brain mask; (3) generating an LNM map for each randomly sampled set, *LNMrandom*; and (4) comparing the observed LNM map with the resulting empirical distribution of the random LNM maps^9, 24, 33^. This produces a voxel- or region-wise LNM z-map and corresponding p-values, which can be optionally followed by e.g. FDR correction or thresholding.

Similar as standard LNM, we can describe these steps in terms of *M* and *C*. The *m* lesion locations represented in the lesion matrix *M* are projected through the normative connectome matrix *C*, leaving *LNMobs = MC*. For each null iteration, the same number of locations is sampled from the *R* possible locations within the brain mask and projected to *C*, yielding *LNMrandom = MrandomC*. Repeating this procedure 5,000 or more times generates an empirical null distribution with mean *μnull* and standard deviation *σnull*, against which *LNMobs* is evaluated, with the null-controlled LNM map conventionally expressed as *LNMz(v) = [LNMobs(v) − μnull(v)]/σnull(v)*, where *μnull(v)* and *σnull(v)* denote the voxel- or region-wise mean and standard deviation of the random LNM across the null-model iterations. **Figure 1** shows the standard LNM pipeline *MC*, extended with the random-lesion null model used by Cui et al. and others^9, 24, 33^.

**Figure 1.**
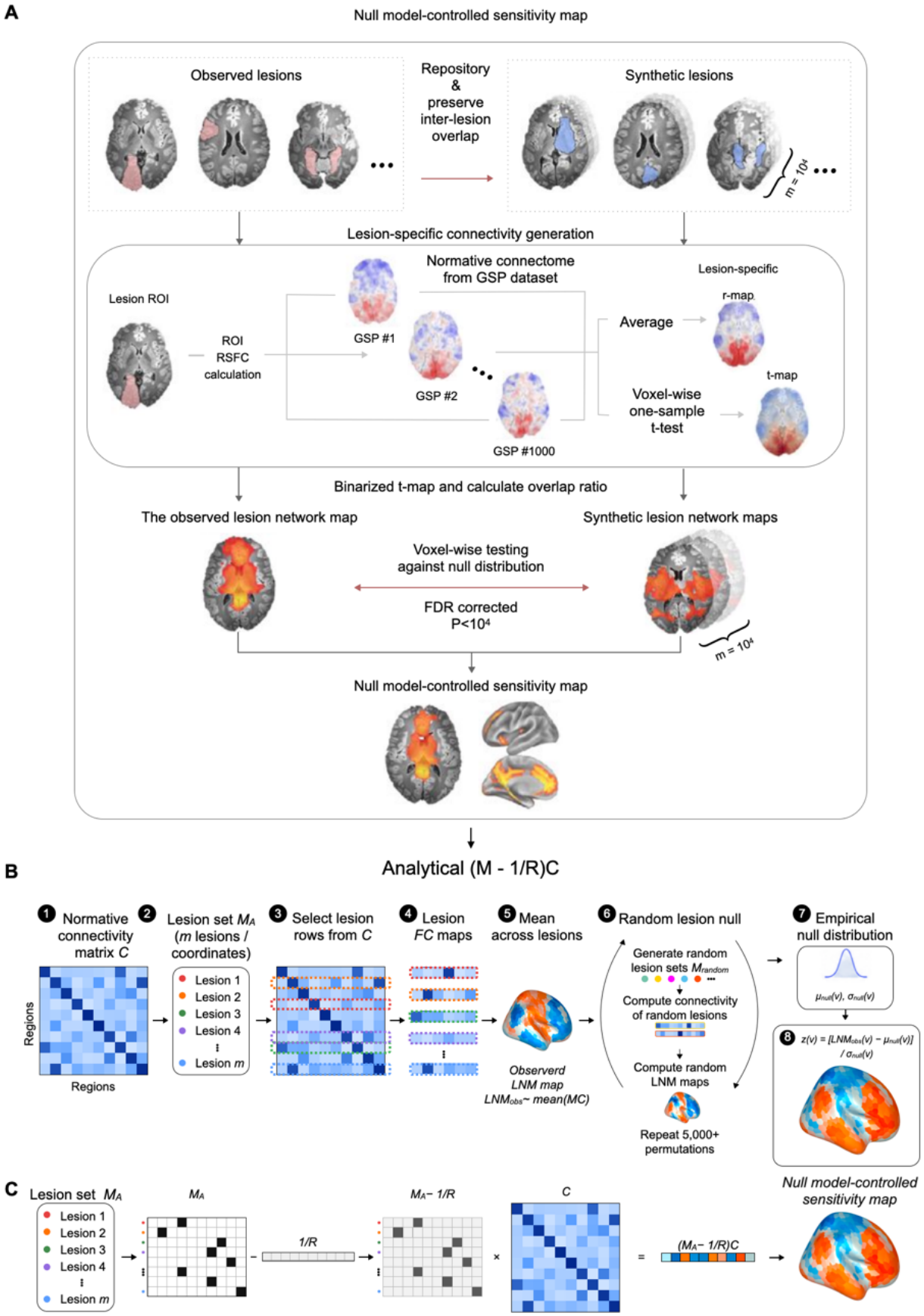
rLNM null-model-controlled sensitivity permutation testing and similar analytical *(M* − *1/R)C* approach. **(A)** Reproduction of the null-model-controlled sensitivity procedure proposed by Cui et al.^24^. An LNM map of patient lesions or coordinates are compared with randomised lesions/coordinates that preserve lesion characteristics^19^. Lesion functional connectivity (fc) maps are derived using the normative GSP1000 connectome^41^ and aggregated into the observed network map. Next, repeating the procedure for 5,000 random lesion sets provides an empirical null distribution against which the observed lesion network map is tested, resulting in the null-model-controlled sensitivity map. **(B)** The same sequence of steps can be summarized in the context of the lesion matrix *M* and a normative connectome matrix *C (1)*, yielding identical results. The normalised lesion matrix *M (2)* describes which rows to select from connectivity matrix *C (3-4)* and averaged into the *LNMobs* map *(5)* by means of *LNMobs∼mean(MC)*. For 5,000 or more iterations *(6)*, random lesion locations are selected, and similarly *LNMrandom∼mean(MrandomC)* maps are formed, with the total set of 5,000 maps used as an empirical null-distribution *(7)*, from which the *LNM*_*RL*_ z-score map for each brain region/voxel *v* is derived *(8)*. **(C)** The same procedure has an analytical notation. The null-controlled LNM map *LNM*_*RL*_ is derived as *LNM*_*obs*_ *− μnull*. The expected mean of the *LNMrandom* maps from the random lesions is *mean(C)*, leaving the null-controlled map as *MC − μC*. Consequently, the contrast of the null-controlled LNM map can be obtained as subtracting the expected value *(1/R)C* from *LNM*_*obs*_, and thus similarly from updating *M = [M − (1/R)1]*, and using this updated matrix in the *LNM ∼ MC* procedure. This leaves the null-controlled map *LNM*_*RL*_ *∼ [M − (1/R)1]C*. Indeed, the full Cui procedure and the analytical formulation leaves almost identical maps (r = 0.96, see also Figure 3). *C*, normative functional connectivity matrix; LNM, Lesion Network Mapping; *M*_*A*_, lesion set of interest.

Similar to standard LNM, we note^23^ that this series of steps has a direct analytical formulation: With uniform random sampling from *R* possible locations in the brain, the expected random lesion vector is *E[Mrandom] = 1/R*, and *μnull* can be derived directly as *mean(C)*, or *μC*. Indeed, the empirically estimated *μnull* is highly similar to the corresponding mean pattern derived directly from *C* (r = 0.998); *σnull* can likewise be closely approximated from the variance structure of *C* (r = 0.997)^23^. Consequently, the main contrast of the null-corrected LNM map can be written directly as

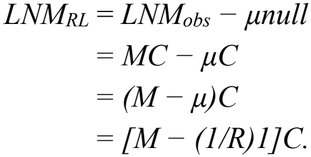

For compactness, we denote this uniform vector as *1/R*, such that *LNM*_*RL*_ *= (M − 1/R)C*.

From the above, it becomes clear that no permutations are needed to obtain the null-controlled sensitivity LNM map. Indeed, the Cui method and a *(M − 1/R)C* implementation produce practically identical LNM maps (r = 0.96, s.d. = 0.04). If desired, values can be scaled by *σC* and a constant *m* to make the overlap exact and on the scale of z-scores.

As such, this step of the rLNM pipeline is almost similar to LNM standard specificity testing, where the network of interest is contrasted against a large (random mixed) group of control lesions^2, 15, 21, 34^, as well as previous reformulations by means of random-coordinate null-models^1, 9, 33^ or case-control contrast LNM^21, 26^. Although an update in the terminology used in LNM procedures can be considered by replacing the term for standard specificity testing with “null-controlled sensitivity testing”, the underlying analytical operations remain the same.

### rLNM cross-symptom specificity testing

The second step proposed by Cui and colleagues extends conventional LNM specificity testing^21^. They build upon the notion of testing LNM contrasts between specific groups, assessing *selective specificity*^24^, contrasting the lesion group of interest to one or more specific control groups and regressing out the common factor shared across the tested conditions.

For three data groups A, B, C, the lesion-seeded fc maps for lesion sets *M*_*A*_, *M*_*B*_, and *M*_*C*_ are selected, and LNM sensitivity maps *LNM*_*A*_, *LNM*_*B*_, and *LNM*_*C*_ are computed by means of the standard LNM approach: *LNM*_*A*_ *∼ M*_*A*_*C, LNM*_*B*_ *∼ M*_*B*_*C*, and *LNM*_*C*_ *∼ M*_*C*_*C* (Figure 1A-B^24^). Next, rLNM proposes combining the maps *LNM*_*A*_, *LNM*_*B*_, and *LNM*_*C*_ into a *cross-symptom map* X, which is regressed out of each standard individual sensitivity map (*LNM*_*A*_, *LNM*_*B*_, *LNM*_*C*_). This regression is argued to produce topologically specific maps^24^, by means of the residuals, *e*_*A*_, *e*_*B*_, and *e*_*C*_ of the regression analysis, with the resulting maps labelled as the rLNM networks. **Figure 2**A illustrates these steps, reproduced from Cui et al.^24^.

**Figure 2.**
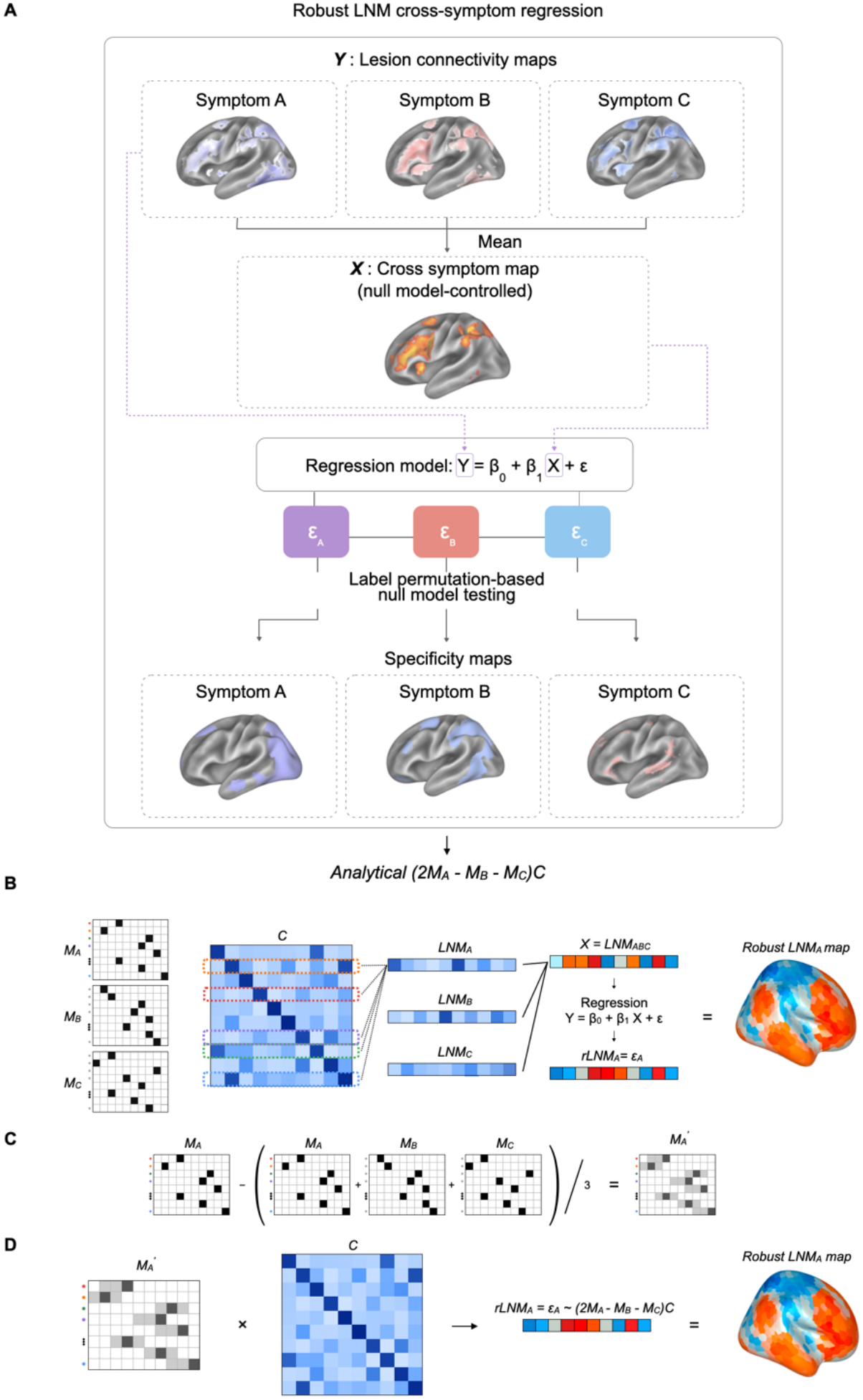
rLNM cross-symptom specificity testing and corresponding analytical reconstruction. **(A)** Reproduction of the cross-symptom specificity framework proposed by *Cui* et al.^24^. *LNM* maps for three symptom groups, A, B, and C are included, with their shared pattern defined by *X = (A + B + C)/3*. The cross-symptom map *X* is regressed from each symptom map, *Y = b0 + b1X + e*, yielding residual maps *e*_*A*_, *e*_*B*_, and *e*_*C*_, which are presented as the *rLNM* maps. **(B)** Schematic representation of the cross-symptom specificity procedure for three symptom groups. Conventional LNM maps are first obtained as *LNM*_*A*_*∼M*_*A*_*C, LNM*_*B*_*∼M*_*B*_*C*, and *LNM*_*C*_*∼M*_*C*_*C*. The cross-symptom map *X=LNM*_*ABC*_, is subsequently used as a predictor of each symptom-specific LNM map, with the regression residual *e*_*A*_ interpreted as the component specific to symptom group A. **(C)** The same operation can be expressed directly at the lesion-matrix level. Removing the mean lesion distribution across the three groups from *M*_*A*_ gives *M’*_*A*_ *= M*_*A*_ *− (M*_*A*_ *+ M*_*B*_ *+ M*_*C*_*)/3 = (2M*_*A*_ *− M*_*B*_ *− M*_*C*_*)/3*. Thus, the procedure contrasts the lesion distribution of group A against those of groups B and C before propagation through the normative connectome. (**D)** Multiplication of the resulting contrast *M’*_*A*_ by the same normative connectivity matrix *C* produces the corresponding residualised LNM pattern, proportional to *(2M*_*A*_ *− M*_*B*_ *− M*_*C*_*)C*. The residualised rLNM map for group A (and similar derivations can be made for group B and group C) therefore remains a linear combination of the original lesion distributions propagated through *C*, rather than introducing an independent source of anatomical information. *M* lesion size and/or lesion set size normalised matrix, *C*, normative functional connectivity matrix; PC, principal component; rLNM, robust Lesion Network Mapping.

We can similarly evaluate these steps in the context of *M* and *C* and find that while the procedure is valuable as a series of complex steps, it again effectively only updates matrix *M*, still leaving the overall approach an *MC* operation. Consequently, the output maps retain repetition and lack specificity when tested across datasets, like standard sensitivity LNM and standard specificity LNM maps.

In rLNM, the *cross-symptom map* X is obtained by averaging the standard LNM maps for conditions A, B, and C, regressing this map out of each condition individually. Cui et al. use the geometric mean and threshold the maps to retain only positive values across symptoms. There is no clear conceptual reason to exclude negative values, as LNM studies commonly interpret both positive and negative effects as meaningful, with negative values equally proposed as potential neurostimulation targets^2, 18, 35^. In what follows, we keep the full range of values and use the conventional arithmetic mean, avoiding arbitrary thresholding^36-39^.

As illustrated in Figure 2A, the rLNM procedure regresses the cross-symptom map X from each symptom map using linear regression analysis, describing *Y = b0 + b1X + e*, where Y is the LNM sensitivity map for symptom A, B, or C individually and X the cross-symptom map. This leaves *LNM*_*A/B/C*_ *= b0 + b1X + e*_*A/B/C*_, where *e*_*A*_, *e*_*B*_, and *e*_*C*_ represent the proposed robust symptom-specific *rLNM*_*A*_, *rLNM*_*B*_, and *rLNM*_*C*_ maps for each condition. When applied to e.g. coordinates related to types of working memory (face, visuospatial and verbal), Cui et al. report a drop in the correlation between the three rLNM maps to around r ≈ 0. This drop is taken as evidence for specificity of the obtained residual rLNM maps, and with that as a solution for the raised limitation of repetition among LNM outcomes.

### The case of 2-group rLNM

When we examine the 2-group rLNM case, the proposed procedure reduces to a specific form. Let the two LNM maps, *LNM*_*A*_ and *LNM*_*B*_, be *A = M*_*A*_*C* and *B = M*_*B*_*C*. Comparing two symptom groups A and B, residualising them against their cross-symptom map *X* is conceptually similar to residualising them against their joint mean, *µAB = (A+B)/2*. The outcome step then becomes mathematically trivial: *e*_*A*_ ≈ *A−µAB = (A−B)/2* and *e*_*B*_ ≈ *B−µAB = (B−A)/2*, with the residuals necessarily perfectly inversely correlated (r = −1). Consequently, a drop in correlation to r = −1 between *rLNM*_*A*_ *= e*_*A*_ and *rLNM*_*B*_ *= e*_*B*_ is not an informative finding, but a direct result of the mathematical procedure performed, with not another outcome possible. When further only positive values are considered in each of the maps—as proposed by rLNM—the resulting specificity maps for symptoms A and B necessarily differ, as all positive values in *eA* are assigned to symptom A while all negative values are assigned to B (and thresholded out), and vice versa for *eB*. Consequently, the thresholded maps will visually appear as highly distinct, while underneath their patterns are highly related (e.g. r = −1). In essence, with two conditions, the rLNM procedure is conceptually analogous to the more transparent two-sample t-test procedure performed in LNM specificity testing, contrasting two groups and assigning the positive values to one group and the negative values to the other group^2, 15, 21^.

### 3-group rLNM

The 3-group comparison is a natural extension of the 2-group rLNM. Let the three LNM maps *LNM*_*A*_, *LNM*_*B*_, *LNM*_*C*_ be *A = M*_*A*_*C, B = M*_*B*_*C*, and *C = M*_*C*_*C*, and their joint mean be *X* = *ABC = (A+B+C)/3*. Residualising each group against this mean gives, for example, for group A, *rLNM*_*A*_ = *e*_*A*_ *= A−ABC = (2A−B−C)/3*. The residual map *e*_*A*_ for group A (*rLNM*_*A*_) is thus a contrast of group A against the average of groups B and C.

This can again be expressed in terms of updating the lesion matrix *M* for each group. With *A = M*_*A*_*C, B = M*_*B*_*C*, and *C = M*_*C*_*C*, and *rLNM*_*A*_ = *e*_*A*_ *∼ A−ABC = (2A−B−C)/3*, the examined contrast is described by *[2M*_*A*_*C −M*_*B*_*C − M*_*C*_*C]/3*, or alternative, *∼ [(2M*_*A*_ *−M*_*B*_ *− M*_*C*_ *)/3]C*. Likewise, *e*_*B*_ *∼ [(2M*_*B*_ *− M*_*A*_ *− M*_*C*_ *)/3]C* and *e*_*C*_ *∼ [(2M*_*C*_ *− M*_*A*_ *− M*_*B*_*)/3]C*. Indeed, adjusting matrix *M*_*A*_ to an updated version *M*_*A*_ *=* (2*M*_*A*_ *− M*_*B*_ *− M*_*C*_; **Figure 2**D) and applying *M*_*A*_*C* with this updated matrix, results in similar rLNM maps as proposed for the full procedure of Cui et al. (r = 0.85, s.d. = 0.11, **Figure 3**); identical maps are obtained when further including fitted beta weights for groups A, B, and C (r = 1.00). Examples are presented at obust.lnmviewer.org.

**Figure 3.**
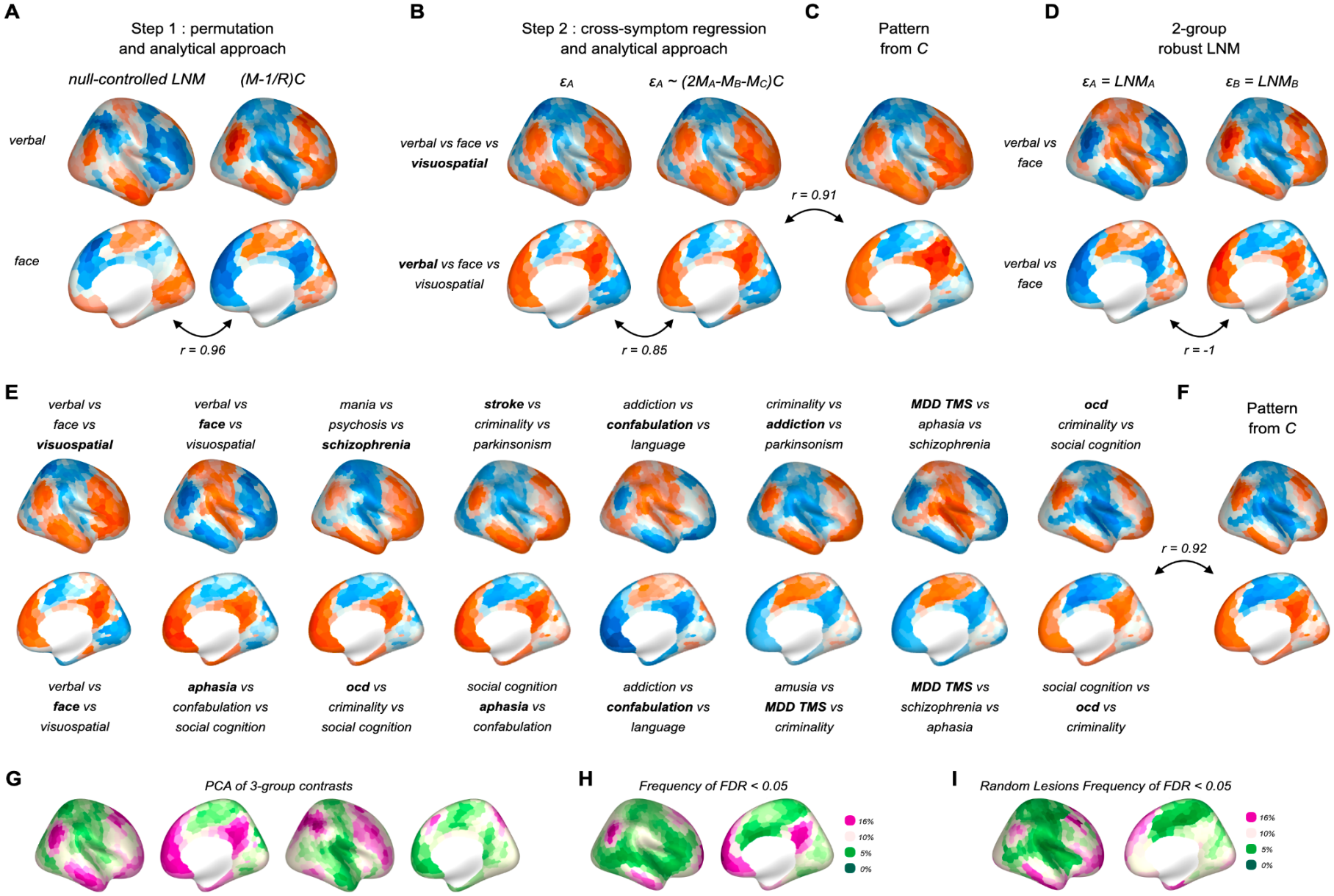
Robust LNM and analytical *(M − 1/R)C* and *(2M*_*A*_ *− M*_*B*_ *− M*_*C*_*)C* leave repetition among LNM outcomes. **(A)** Null-controlled LNM sensitivity procedure and analytical expression *(M − 1/R)C* results in similar maps. Examples shown for verbal and face working memory. **(B)** The second step of the proposed robust LNM pipeline, regressing out the cross-symptom map among three examined groups A, B, C from each of the LNM maps, with the residual map *e*_*A*_, *e*_*B*_, *e*_*C*_ describing the robust LNM maps, can be also reduced to an analytical formulation LNM *e*_*A*_ *∼ (2M*_*A*_ − *M*_*B*_ − *M*_*C*_). Example maps are shown for a 3-group comparison of verbal, face, and visuospatial working memory, with visuospatial and verbal memory maps shown in B for both the full permutation as well as analytical formulation (average r = 0.85 across 1,000 3-group comparisons). The analytical notation shows that the procedure only updates matrix *M*, leaving the core procedure similar to *∼MC*. **(C)** The null-controlled sensitivity map and the robust LNM map *rLNM*_*A*_, or *e*_*A*_, remain patterns of *C*. Panel shows a regression analysis of the rLNM maps presented in panel B against the linear regression model (see text), showing that rLNM maps are similarly determined by a small number of basic standard patterns of *C* (subcortical, cortical degree, 4 modules, 3 gradients) **(D)** The proposed *rLNM* procedure has a special form when performed in a 2-group design. Here, regressing the cross-symptom map X between group A and B out of both *LNM*_*A*_ and *LNM*_*B*_ leaves their residuals *e*_*A*_ and *e*_*B*_, and with that the robust r*LNM*_*A*_ and *rLNM*_*B*_ maps, trivially maximally different (they are exactly each other’s opposite, r = −1). **(E)** When rLNM is performed in a 3-group design as discussed by Cui et al. the maps are also naturally distinct, but nevertheless remain patterns of *C* and thus remain repetitive when compared with other 3-group *rLNM* maps. The panel shows several contrasts within the 3-group rLNM design, with the shown panel highlighted in bold. **(F)** Similar to the standard sensitivity, specificity and null model LNM maps^21, 33^, also rLNM maps remain a pattern derived from *C*. Panel shows an example of a standard pattern of *C* derived by means of the simple regression model. **(G)** Principal component analysis (PCA) across 3-group rLNM contrasts reveals strong similarity among maps across conditions, with the first five PCs explaining 83% of the variance (28 lesion datasets). Panels show the first two PCs. **(H)** Of all 3-group contrasts, 55.7% contain at least one significant region (p_FDR_ < 0.05), with the spatial frequency of significant findings showing spatial similarity with PC1 of connectivity matrix *C* (r = 0.48). **(I)** Repeating the analysis using spatially randomised lesion sets (10,000 3-group contrasts) reveals a similar spatial pattern of significant findings, with 47.5% of maps containing at least one significant region (p_FDR_ < 0.05) despite randomisation. The spatial distribution of frequency of significant findings correlates with PC1 of C (p_FDR_ < 0.05, r = 0.66).

In the 2-group case, *e*_*A*_ *= −e*_*B*_. Similarly, in 3-group rLNM, *e*_*A*_, *e*_*B*_, *e*_*C*_ follow the constraint *e*_*A*_ *+ e*_*B*_ *+ e*_*C*_ *=* 0 at each brain region/voxel, with any positive deviation in one group balanced by negative deviations across one or both other groups. When next the positive residuals are thresholded and retained per map as proposed by Cui et al., each resulting map highlights locations where that group exceeds the average of the other two. A low correlation between *e*_*A*_, *e*_*B*_, *e*_*C*_ after the rLNM procedure (reported by Cui et al. as r = 0) can thus not be taken as evidence for specificity of the retained rLNM maps, as a low correlation follows directly from the mathematical steps performed.

Label shuffling is used to obtain a permutation-based reference distribution. This permutation testing changes the reference against which effects are tested, but it does not change the main contrast examined, which in rLNM are the described residual maps *e*_*A*_, *e*_*B*_, *e*_*C*_ . Once the group maps are residualised against their shared mean, they represent constrained contrasts among A, B, and C. In the 2-group case, this is especially transparent: Residualisation reduces exactly to the A*−*B contrast, such that label shuffling becomes a permutation-based 2-group comparison, with resulting z-values analogous to the standard two-sample test used in LNM specificity testing^22^. With three groups, permutation similarly tests how unusual each group-versus-others contrast is under random reassignment of group labels. It therefore changes the statistical reference used to evaluate the contrast, and hence the effects that survive further statistical thresholding, but not per se the rLNM maps being evaluated, which remain derived solely from matrix *C*.

### Empirical testing

We performed the rLNM approach for 2-group and 3-group contrasts on multiple lesion and coordinate datasets, including rLNM maps for coordinates related to face, visuospatial, and verbal working memory, as described by Cui et al.^24^. We complemented this data with 67 other lesion sets for e.g. addiction, aphasia, epilepsy, etcetera, as symptom group A, B, and C, respectively (see for a full description of the data^14^).

We derived similar null-controlled LNM maps as computed by the full permutation approach (10,000 permutations) using *rLNM ∼ (M − 1/R)C* (r = 0.96, s.d. = 0.04, **Figure 3A**). Also rLNM maps derived in step 2 of the rLNM procedure were equally derived by means of simply adjusting *M′*_*A*_ to *2M*_*A*_ *− M*_*B*_ *− M*_*C*_ and applying *M′*_*A*_*C* (r = 0.85, s.d. = 0.14; 1,000 random 3-group lesion set combinations, **Figure 3**B).

We next examined the rLNM maps *e*_*A*_, *e*_*B*_ and *e*_*C*_ for coordinates related to face, visuospatial, and verbal working memory in more detail. Comparing them against the standard patterns of *C* (linear regression model^14^) revealed that rLNM maps are similarly described by means of the standard factors of *C* (R^2^ = 0.79, 0.89, 0.76), similar as reported for the standard LNM maps^14^. To examine this more generally, we repeated the rLNM procedure for 10,000 combinations of 3 groups sampled from the total 67 LNM lesion sets. This confirmed that *e*_*A*_, *e*_*B*_, and *e*_*C*_ rLNM maps strongly link to *C*’s standard properties, with *C*’s standard factors explaining on average 83% (s.d. = 0.14) of the spatial variance in rLNM maps across the 3-group combinations.

Although the rLNM procedure^24^ is designed to mitigate the disease-specificity problem of LNM^22^, it becomes clear that the resulting maps remain highly repetitive when we compare them against the rLNM maps of other contrast groups. For example, rLNM returns a similar ‘visuospatial rLNM network’ from a 3-group comparison, when we for example, examine a mania-psychosis-schizophrenia contrast (r = 0.82, **Figure 3E**) or a creativity-criminality-OCD comparison (r = 0.90, **Figure 3**E), conditions that are unrelated to the working-memory network examined. This persistent repetition among rLNM maps is illustrated by a principal component analysis across 10,000 derived rLNM 3-group maps, showing that a limited number of standard patterns determine rLNM’s outcomes: 5 simple patterns across the tested 3-group rLNM maps explain as much as 83% of their variance, with the principal components naturally aligned with the PCs of matrix *C* (82%, **Figure 3**G).

### Significant regions

We examined the number and spatial distribution of significant regions across the 3-group lesion-set contrasts when maps are further tested. We computed ∼10,000 contrasts (9,828 contrasts across 28 randomly sampled lesion sets, together describing 3,171 lesions), of which 5,477 (55.7%) contained at least one significant region (FDR *q* < 0.05). The spatial distribution of these positive findings across brain regions correlated with the first principal component (PC1) of the connectivity matrix *C* (r = 0.48), consistent with previous observations of LNM specificity analyses to report overlap in published peak locations across literature^22^. We repeated the analysis using random lesion sets. Lesion locations were randomised across the cortex, preserving spatial overlap by spinning the lesions around the cortex^22^ and we sampled a similar 10,000 3-group contrasts from these randomised sets. Despite randomisation of coordinates/lesions, a similar 47.5% of resulting rLNM maps showed regions surviving FDR correction (*q* < 0.05, **Figure 3**I), effects overlapping as those reported for standard LNM^14^.

## Discussion

The proposed rLNM framework^24^ combines several statistical concepts within LNM. While forming a series of elaborate steps removing one shared pattern in LNM maps, the proposed procedure does not change the source of the information being tested, which remains the normative connectivity matrix *C* and a *MC* operation. The raised concern about LNM is therefore not that two or three maps cannot be statistically distinguished, but that when applied to multiple conditions, maps are repetitive and overlapping. Adding new statistical layers may further change the contrast, re-weight the signal, or introduce a new reference distribution, but they do not remove LNM’s core underlying methodological constraint to *C*, leaving persistent repetition when applied to multiple conditions and datasets.

The relevance of adding methodological steps to LNM deserves careful consideration, as increasingly complex workflows may ultimately obscure an underlying methodological dependency. Presenting elaborate procedures as solutions without extensive testing or evaluation across multiple datasets, risks overstating what is being achieved. Designing a new reference condition to which LNM maps can be tested^26, 33^ may recalibrate inference, and as such be helpful in the statistical benchmarking of LNM maps, but it does not expand the space of recoverable patterns in *C*. Instead, one could argue that relabeling previous steps or adding unnecessary complexity may make the line of reasoning harder to follow, while leaving the core procedure intact.

Cui and colleagues propose rLNM as a new variant to the LNM framework. rLNM may remove a shared connectome-derived pattern, such as degree or a 3-group cross-symptom map X, and may therefore be useful for reducing the influence of this component in LNM maps. However, rLNM does not generate fundamentally new types of LNM maps. Instead, it primarily shifts the resulting network toward other repetitive patterns within *C*, without solving the inherent constraints imposed by *C*. Repetition and overlap among unrelated conditions remain an expected outcome of LNM networks, including those derived by robust reformulations.

## Methods

### Lesion sets

Lesion and coordinate datasets were obtained from previously assembled collections of published psychiatric and neurological lesion network mapping (LNM) studies, as described in ^14^. 67 lesion and coordinate sets were included to examine the similarity and anatomical specificity of LNM (rLNM) maps^14^.

### Lesion network mapping

Voxel-wise analyses were performed using Lead-DBS v3.1^40^ with MATLAB R2024b, SPM12, and the GSP1000 normative functional connectome^41^. Lesions were analysed in MNI152 space using the lead_mapper interface and cs_fmri_conseed_seed_tc.m function. Atlas-based LNM used the Yeo-Schaefer1000 cortical atlas^42^ and Melbourne54 subcortical atlas^43^, selecting connectivity profiles corresponding to lesion locations from the normative connectivity matrix *C*.

### Robust lesion network mapping

Robust LNM (rLNM) was evaluated using 2- and 3-group contrasts following the framework proposed by Cui et al.^24^. Conventional LNM maps were obtained for each lesion set using the normative functional connectome. The cross-symptom map X was defined as the arithmetic mean of the individual LNM maps, retaining both positive and negative connectivity values. The cross-symptom map was subsequently regressed from each individual LNM map, with the residuals representing the corresponding rLNM maps (see main text for details).

### Analytical reconstruction of rLNM

The null-controlled sensitivity procedure was compared with its analytical formulation, *(M − 1/R)C*, where *M* represents the normalised lesion distribution, *R* the number of possible lesion locations, and *C* the normative functional connectome. Similarly, 3-group rLNM contrasts were reconstructed using the lesion-matrix formulation *(2M*_*A*_ *− M*_*B*_ *− M*_*C*_*)C*, corresponding to subtraction of the cross-symptom mean. An exact reconstruction incorporating the fitted regression coefficients was additionally evaluated. Correspondence between the analytical reconstructions and the original procedures was assessed using Pearson spatial correlations.

### Correspondence with normative connectivity patterns

The contribution of normative connectome properties to rLNM maps was examined using multiple linear regression, following the approach described previously^14^. Predictors included cortical and subcortical connectivity degree, four functional modules, and three principal connectivity gradients derived from the GSP1000 group-averaged connectome matrix *C*. The proportion of spatial variance explained by these normative patterns was quantified using the coefficient of determination (R^2^). Principal component analysis was additionally performed across rLNM maps to quantify their dimensionality. Repetition across conditions was assessed using pairwise Pearson spatial correlations.

### Statistical testing and spatial null models

Statistical significance was evaluated across ∼10,000 3-group combinations (9,828 contrasts derived from 3,276 3-group combinations of 28 publicly available lesion datasets^14^). Group labels were permuted to generate reference distributions for the observed rLNM contrasts, and statistical significance was assessed at the regional level using false discovery rate correction (*q* < 0.05). The proportion of contrasts containing at least one significant region was calculated, together with the spatial frequency of significant findings across the brain. This spatial distribution was compared with the first principal component of the normative connectivity matrix using Pearson correlation. To examine the same effect for random lesions, and show its independence of real lesion locations, the analysis was repeated using spatially randomised lesion sets, preserving the spatial overlap of the original lesions^22^. A similar total of 10,000 3-group contrasts were examined using these randomised datasets, and the proportion containing at least one significant region was compared with that observed for the original lesion sets.

